# Novel immunotherapy for multiple solid cancers using an Anti-HVEM blocking monoclonal antibody

**DOI:** 10.1101/2024.12.16.627913

**Authors:** Gilli Galore-Haskel, Efrat Merhavi-Shoham, Mika Shapiro, Roni Bareli, Naama Dror, Sivan Seliktar-Ofir, Katerina Shamalov, Adva Levy-Barda, Goni Hout-Siloni, Jacob Schachter, Eran Sadot, Ram Eitan, Iris Barshack, Shay Golan, Yael Feferman, Effi Yeoshoua, Saeb Eyadat, Eyal Greenberg, Gal Markel

## Abstract

**INTRODUCTION:** Immune checkpoint inhibitors (ICIs) have revolutionized cancer treatment, yet their efficacy remains limited. Therefore, there is a clear need for new anti-tumor agents. HVEM (Herpes Virus Entry Mediator) plays a regulatory role in immunity, making it a promising cancer therapeutic target.

**EXPERIMENTAL DESIGN:** We have developed Anti-4CB1, a fully human HVEM-BTLA and HVEM-CD160 blocking mAb and tested its anti-tumor activity in various *in-vitro, ex-vivo* and *in-vivo* models, alone or in combination with Anti-PD1. Finally, we analyzed HVEM expression in serum samples and melanoma tumor tissues and its correlation with HVEM and PD1 blockade treatment response.

**RESULTS:** *In-vitro* assays demonstrated enhanced melanoma cell killing by autologous TILs in the presence of Anti-4CB1. In addition, Anti-4CB1 significantly increased cytotoxicity by up to 233% in a variety of *ex-vivo* cancer tissue samples of different indications. Notably, Anti-4CB1 demonstrated effectiveness in samples where Anti-PD1 was ineffective. In addition, significant anti-tumor activity at 10 mg/kg in combination with Anti-PD1 (TGI 95% p=0.0001) or at 25 mg/kg as monotherapy (TGI 50% p=0.0096) was demonstrated in a colon carcinoma transgenic mice model. Moreover, significant anti-tumor activity was observed in a mouse ImmunoGraft model (TGI 53% p=0.0102 as monotherapy). Finally, we demonstrated that HVEM expression correlated with response to Anti-HVEM and Anti-PD1 treatments.

**DISCUSSION:** We describe the development of Anti-4CB1, a fully human anti-HVEM mAb, blocking HVEM-BTLA and HVEM-CD160 human interactions and enhancing lymphocytes cytotoxicity against various cancer cells *ex-vivo*. *In-vivo,* Anti-4CB1 showed promising anti-tumor effects, particularly in combination with Anti-PD1, suggesting its therapeutic potential. Finally, HVEM expression may serve as a predictive marker for Anti-HVEM and Anti-PD1 treatments response, offering potential diagnostic utility in patient selection for these immunotherapies. These results support Anti-4CB1 potential as an anti-cancer therapeutic agent.

**Translational Relevance:** The discovery of immune checkpoint inhibitors (ICIs), marks a significant breakthrough in cancer therapy. However, despite the remarkable success of current ICIs targeting CTLA4 and PD1/PD-L1 pathways, a significant proportion of patients fail to respond or develop resistance. Therefore, the search for novel ICIs is crucial for advancing cancer immunotherapy. Anti-HVEM emerged as a promising candidate, as we demonstrate in *ex-vivo* and *in-vivo* models, its ability to block the HVEM-BTLA interaction, resulting in enhanced antitumor immune responses in solid tumors, alone or in combination with existing ICIs, to improve treatment outcomes. Moreover, the correlation between HVEM expression and treatment responses highlight its potential as an attractive target for biomarker-driven therapies. Investigating HVEM as an ICI offers a promising path to expand therapeutic options, personalize treatment approaches, address the evolving challenges in cancer immunotherapy and improve patient outcomes.

## Introduction

Immunotherapy is a field of immunology that aims to identify treatments for diseases through induction, enhancement or suppression of an immune response. Activating immunotherapies, which are designed to boost immune responses, are at the forefront of cancer treatment. Among these, immune checkpoint blockade therapy primes the immune system to attack tumor cells and has been successfully used to treat various cancer types including advanced non-small cell lung cancers (NSCLC), metastatic melanoma, advanced renal cell carcinoma (RCC), metastatic urothelial carcinoma, head and neck squamous cell carcinoma (HNSCC), MSI-high tumors, Merkel cell carcinoma, squamous cell carcinoma, hepatocellular carcinoma, cervical carcinoma and many others (1). A few such drugs have shown significant promise and have received FDA approval, including antibodies targeting the immune checkpoints programmed cell death protein 1 receptor (PD1), lymphocyte activation protein 3 (LAG3) in combination with PD1 blockade, programmed cell death protein 1 ligand (PD-L1) and cytotoxic T lymphocyte-associated protein 4 (CTLA4) (2–17). While these treatments have extended progression-free survival and improved overall survival for some patients, the response rate is still not optimal. Therefore, there is a pressing need to identify new anti-tumor immune activating agents.

HVEM (also known as TNFRSF14), a member of the tumor necrosis factor (TNF) receptor family, was originally identified as the entry route for herpes simplex virus (HSV) (18). It is expressed on the surface of T cells, B cells and other hematopoietic cells, as well as on endothelial and epithelial cells (19,20). HVEM is unique among TNF receptors as it binds both TNF superfamily molecules such as LIGHT (also known as TNFSF14) and soluble TNFβ as well as immunoglobulin molecules such as BTLA (B- and T-lymphocyte attenuator) and CD160, which are primarily expressed on hematopoietic cells (20). The HVEM-BTLA interaction was the first to illustrate crosstalk between these two receptor families. HVEM can function either as a signaling receptor or as a ligand for its interacting molecules, enabling bidirectional signaling for immune regulation under different contexts (21,22). The extracellular domain of HVEM consists of four Cysteine Rich Domains (CRDs) (23); LIGHT and TNFβ bind to CRD2 and CRD3 (24) while BTLA, CD160 and gD bind to CRD1 on the opposite surface of the molecule (25). The cellular context (*cis* or *trans*) and the specific HVEM ligand determine whether the cellular response is activating or inhibitory. In the *trans* configuration, signaling through HVEM turns on NF-κB dependent survival genes, whereas BTLA signaling is inhibitory through SHP1/2 phosphatases. In the *cis* configuration of HVEM-BTLA, BTLA suppresses HVEM signaling of NF-κB by all its ligands in *trans*. Only the LIGHT membrane form can activate HVEM when expressed in the *cis*-complex with BTLA (26). HVEM delivers coinhibitory signals through CD160 in T cells and NKT cells and blocking this interaction has been shown to reverse T-cell exhaustion and enhance production of Granzyme B in virus-specific effector CD8+ T cells (22,27). Naive T cells express high levels of HVEM, which decrease during activation, with high levels being re-expressed at the end of the activation phase (28).

Abnormal expression of HVEM has been reported in different cancer types, both lymphoid malignancies as well as solid tumors. For example, HVEM overexpression in colorectal cancer tissues correlated with tumor invasion and pathological stage, and resulted in significantly poorer prognosis for patients with high HVEM expression compared to those with lower expression (29). Similarly, HVEM expression was associated with aggressive tumor features in invasive breast cancer (30) and significantly correlated with postoperative recurrence and survival in hepatocellular carcinoma (31). HVEM is expressed in the majority of cultured melanoma cell lines and metastatic melanoma samples. Of importance, the interaction of BTLA on tumor specific T cells and HVEM on melanoma cells resulted in T cell inhibition (32).

Although HVEM-BTLA axis has already been suggested as potential therapeutic target for cancer, to the best of our knowledge, only one pre-clinical work has been published recently (33,34), while no clinical trials targeting HVEM for cancer therapy have been initiated so far.

Here, we describe the development and some preclinical characterization of a fully human anti-HVEM mAb (named Anti-4CB1) capable of specifically blocking human HVEM-BTLA and HVEM-CD160 interactions. Anti-4CB1 activates TILs and PBMCs leading to enhanced cytotoxicity towards various cancer cells *ex-vivo*. In addition, *in-vivo* studies in humanized mice using PDXs and in hBTLA and hBTLA/hHVEM transgenic immuno-competent mice model, show anti-tumor effect of the mAb alone or in combination with Anti-PD1, suggesting Anti-4CB1 as a potential therapeutic agent for melanoma, colon and other cancers. Lastly, HVEM expression was found to be a predictive marker for Anti-HVEM and Anti-PD1 treatments and may be used diagnostically to determine a patient population that is more likely to respond to these immunotherapies.

## Materials and Methods

### Cells, blood and serum samples

Human bladder, colon, endometrial, gastric, hepatocellular, lung, melanoma, ovarian, pancreatic and renal cancer tissues were obtained from surgically excised specimens at Sheba Medical Center and Rabin Medical Center. Specimens were received with autologous fresh PBMCs from the cancer patients. Enzymatic digest and *ex-vivo* expansion of TILs was performed as previously described (35). When available, serum samples from these cancer patients were obtained along with serum samples from healthy donors. Jurkat E6.1 and A549 cell line were purchased from ATCC. Melanoma cells used for *in-vitro* experiments and Jurkat E6.1 cells were maintained with RPMI-1640 medium with 10% FBS and 1 mM sodium pyruvate (Sartorius, #03-042-1B). A549 cell line was maintained in DMEM with 10% FBS. All mediums contained 100 U/mL penicillin-streptomycin (Sartorius, #03-031-1B), 25 mmol/l HEPES pH 7.2 (IMBH, # L0180-100) and 2 mM L-glutamin (Sartorius, #03-020-1B). Primary TILs bulk and PBMCs cultures were cultured in RPMI-1640 supplemented with 10% heat inactivated human serum (Valley, #HP1022), 100 U/mL penicillin-streptomycin, 50 μM 2-mercaptoethanol (Gibco, #31350010), 25 mmol/l HEPES pH 7.2 and 6,000 IU/mL or 600 IU/mL IL-2, respectively (Proleukin, Novartis). Macrophages were cultured in RPMI-1640 supplemented with 10% heat inactivated human serum, 100 U/mL penicillin-streptomycin, 10 mmol/l HEPES pH 7.2 and 1 mM sodium pyruvate. CHO-K1 cells (ECACC, #85051005) were maintained in F-12 Nutrient Mixture (Ham’s) with 10% FBS. Cell density was maintained between 0.5×10^6^-2×10^6^/mL. Human peripheral blood was obtained from healthy donors from MDA Israel. Monocytes were isolated using RosetteSep™ Human Monocyte Enrichment Cocktail (Stemcell Technologies, #15068) and Ficoll/Hypaque cushion, according to manufacturer’s instructions. Then monocytes were maturated into macrophages M1 by incubating with macrophages media containing 50 ng/mL M-CSF (Peprotech, #300-25-10) for 5 days following incubation of 2 days with media containing IFNγ 20 ng/mL (Biotest, #285-IF-100) and LPS 100 ng/mL (Sigma-Aldrich, #L2630). Differentiation of monocytes into M1 macrophages was confirmed by flow cytometry. All studies were performed under approved protocols with written informed consent from patients and in line with the principles of the Declaration of Helsinki.

### Generation of stable expression cell systems

CHO-K1 cells were seeded in 6-well plates (2x10^5^ cells/well) and incubated overnight at 37°C. Following incubation, transfection was performed using jetPRIME transfection reagent (Polyplus, #114-01), according to manufacturer’s instructions. Empty pcDNA3.1 vector (control), human HVEM (hHVEM), mouse HVEM (mHVEM) and cynomolgus HVEM (cHVEM) constructs subcloned into pcDNA3.1 vector were purchased from Genscript. Selection with 1 mg/mL of G418 antibiotics (Mercury, #345810) was performed two days post-transfection and cells were routinely cultured with addition of antibiotics. In addition, we generated CHO-K1 HVEM/TCR activator cells - CHO-K1 cells overexpressing human HVEM with the TCR Activator Lentivirus (BPS Bioscience, #79894-CP). The Lentivirus contains a gene for a membrane-bound, engineered T cell receptor (TCR) activator driven by a CMV promoter. For generating nuclear RFP melanoma cells, cells were seeded in 96-well flat plates (5x10^3^ cells/well) and incubated overnight at 37°C. Then, the medium was aspirated and replaced with RPMI-1640 mediumcontaining Polybrene (Sigma Aldrich, #TR-1003-G) and IncuCyte NucLight Lentivirus Reagent (Essen BioScience, #4625) to reach a MOI of 3. Cells were incubated overnight at 37°C, after which transduction media was replaced with fresh media. Transfection efficiency was assessed 48 hours after transduction by Incucyte live-cell analysis system (Sartorius, Germany).

For generating recombinant Jurkat T cell expressing eGFP gene under the control of NFAT response elements, Jurkat E6.1 cells were transduced with NFAT eGFP reporter lentivirus (BPS Bioscience, #79922). The lentivirus vector contains an enhanced GFP gene driven by the NFAT response element located upstream of the minimal TATA. After transduction, activation of the NFAT signaling pathway in the cells was monitored by examining eGFP expression by Incucyte live-cell analysis system and flow cytometry. For constitutive expression of human BTLA (hBTLA) or human CD160 (hCD160), Jurkat NFAT eGFP cells were transduced with CD272 (BTLA) (NM_181780) Human Tagged ORF Clone Lentiviral Particles (Origene, #RC219458L3V) to generate Jurkat BTLA/NFAT-eGFP cells or CD160 (NM_007053) Human Tagged ORF Clone Lentiviral Particles (Origene, #RC204886L3V), to generate Jurkat CD160/NFAT-eGFP cells respectively. Successful expression of human BTLA and CD160 was confirmed by flow cytometry.

### Generation of scFv against HVEM by phage display

The generation of scFv was performed by RxBiosciences (Gaithersburg, MD, USA) Naïve human scFv library (>10^11^ clones) was incubated with recombinant tags in order to subtract tag specific clones and with control CHO-K1 cells in order to subtract CHO-K1 cell binding clones. Then, phages were subjected to three biopanning rounds through incubation with CHO-K1 cells overexpressing human HVEM. Clones were picked, rescued with the help of helper-phage and screened by phage ELISA against human (R&D Systems, #356-HV), mouse (R&D Systems, #2516-HV) and cynomolgus (R&D Systems, #9197-HV) recombinant HVEM proteins. Clones were sequenced using Sanger’s method and DNA sequences were analysed by CLC Bio software (Qiagen). Positive unique clones were sub-cloned into pET25b protein expression vector and recombinant DNA was transformed in BL21 cells. Lysates were prepared from 200 mL of induced culture. Bacterial cells were pelleted, washed with PBS and lysed by French press. Insoluble debris was removed by high-speed centrifugation and the lysates containing the soluble proteins, including the scFvs, were collected. The scFvs were cloned with HIS tags.

### Flow cytometry

Staining for extracellular and intracellular proteins was performed according to standard protocols, as previously described (36). Gating of cells was performed using FSC vs. SSC. Background fluorescence intensity was set by isotype control or secondary antibody only stained samples. All experiments were performed using a CytoFLEX instrument (Beckman Coulter) and data was analyzed using FCS Express software (De Novo Software) and GraphPad Prism.

### Antibodies used

Anti-4CB1 proprietary anti-HVEM mAb; Human IgG4 isotype control (Sino Biological Inc. #HG4K); HIS Tag APC-conjugated Antibody (R&D Systems, #IC050A); Human HVEM/TNFRSF14 polyclonal antibody (R&D Systems, #AF356); Alexa Fluor 488 Rabbit anti-Goat IgG (Invitrogen, #A11078); Anti-melanoma antibody (Abcam, #ab732); PE/Cyanine7 anti-human CD86 Antibody (BioLegend, #374209); AlexaFlour 647 anti-human CD80 Antibody (BioLegend, #305215); Alexa Fluor 488 anti-human CD14 Antibody (BioLegend, #325610).

### Binding and blocking ELISA

For binding ELISA, plates were coated with recombinant proteins for overnight at 4°C. Plates were washed with PBS and blocked with 2% BSA (Sigma Aldrich, #A3059) for one hour at room temperature to eliminate non-specific binding. Tested antibodies, diluted in blocking buffer, were added to the plates and incubated for one hour at room temperature. Plates were washed and incubated for one hour at room temperature with detection antibody, followed by washes and addition of peroxidase anti-human FC for 30 minutes at room temperature. Finally, plates were washed and TMB (Mercury, #ES001) followed by stop solution 0.5M H_2_SO_4_ (Sigma Aldrich, #258105) was added. For blocking ELISA, after addition of tested antibodies, plates were washed, and recombinant proteins were added to plates and incubated for two hours at room temperature. Plates were washed and incubated for one hour at room temperature with antibodies detecting the recombinant proteins. Plates were washed and streptavidin peroxidase or peroxidase specific antibody was added for 30 minutes at room temperature followed by the steps mentioned above. Absorbance was read at 450nM and 570nM.

### Recombinant proteins and antibodies used

For human HVEM binding ELISA: Human HVEM-His tag Protein (ACRO, #HVM-H52E9), peroxidase anti-human FC (Jackson, #109-035-098).

For mouse HVEM binding ELISA: mouse HVEM HIS+Fc tag, (Sino Biologicals, #10567-M03H); Peroxidase Streptavidin, (Jackson, #016-030-084).

For human HVEM-BTLA blocking ELISA: human HVEM HIS tag, (Sino Biologicals, #10334-H08H); hBTLA-Fc&AVI protein (Sino Biological, #11895-H41H-B); Peroxidase Streptavidin, (Jackson, #016-030-084).

For human HVEM-LIGHT blocking ELISA: human HVEM HIS tag, (Sino Biologicals, #10334-H08H); Human LIGHT Protein HIS Tag, (Sino Biological, #10386-H07H); Human LIGHT/TNFSF14 Antibody, (R&D Systems, #AF664); Peroxidase-AffiniPure Bovine Anti-Goat IgG, (Jackson, #805-035-180).

For human HVEM-CD160 blocking ELISA: human HVEM HIS tag (Sino Biologicals, #10334-H08H); Human CD160 HIS tag Biotin labeled protein (BPS BioScience, #71135-1); Peroxidase Streptavidin (Jackson, #016-030-084).

For mouse HVEM-BTLA blocking ELISA: mouse BTLA Fc tag, (Sino Biologicals, #51060-M02H); mouse HVEM HIS tag, (Sino Biologicals, #10567-M08H); BTLA/CD272 Biotin Antibody (Novus, #HMBT-6B2); Peroxidase Streptavidin, (Jackson, #016-030-084)

### Cytotoxicity, 41BB and CD107a expression and apoptosis by time lapse microscopy

Nuclear RFP-stained melanoma cells were seeded with IFNγ (R&D Systems, #285-IF-100) in 96-well plates and incubated overnight at 37°C and 5% CO_2_ to allow them to attach. Tested antibodies were added for one hour pre-incubation with cells. Then, given amounts of TILs, or medium only, were added to the cells. In *ex-vivo* experiments using fresh tissues, digested tumor cells were seeded together with autologous PBMCs and tested antibodies with no pre-incubation. For cytotoxicity and apoptosis assays, CellEvent Caspase-3/7 Green Detection Reagent (Invitrogen, #C10423) was added too and Caspase 3/7 positive events, reflecting specific killing of cancer cells, were counted. For 41BB expression, Incucyte FabFlour-488-labeled (Satorius, #4745) anti-41BB antibody (BioLegend, #309802) was added together with TILs to melanoma cells. 41BB positive events, reflecting T-cells activation, were counted. For CD107a expression, TILs or medium were added to the melanoma cells together with conjugated CD107a antibody (BioLegend, #328620) and positive events were counted. All co-cultured plates were placed in the Incucyte live-cell analysis system (Sartorius, Germany) and scanned every hour.

### Cell-based Co-inhibitory Bioassay

BTLA/NFAT: CHO-K1 HVEM/TCR activator cells were seeded in 96-flat plate. Next, cells were incubated for one hour at 37°C with 0.2-20 μg/mL of Anti-4CB1 or isotype control. Then, Jurkat BTLA/NFAT-eGFP cells or medium were added to CHO-K1 cells. eGFP reporter activity was tested by placing the plates in the Incucyte live-cell analysis system (Sartorius, Germany) and scanned every hour.

CD160/NFAT: 96-flat plate was coated with 1000 ng/mL OKT3 diluted in PBS. After overnight incubation and washes, Jurkat CD160/NFAT-eGFP cells were seeded. Dynabeads M-280 streptavidin (Invitrogen, #11205D) were incubated with or without HVEM HIS tag protein (Sino Biologicals, #10334-H08H) for 30 minutes with shaking, washed and incubated for one hour at 37°C with 20 μg/mL of Anti-4CB1 or isotype control. After incubation, coated Dynabeads were washed and added to the Jurkat CD160/NFAT-eGFP cells. eGFP reporter activity was tested by placing the plate in the Incucyte live-cell analysis system (Sartorius, Germany) and scanned every hour.

### Phagocytosis assay

M1 macrophages were counted and 1x10^4^ cells/well were seeded in 96-well plate. At the same time, 100 ng/mL IFNγ was added to A549 cells, and both were incubated overnight at 37°C and 5% CO_2_. Then, A549 cells were harvested and labeled by IncuCyte® pHrodo® Red Cell Labeling Kit for Phagocytosis (Satorius, #4649), according to the manufacturer’s protocol. Tested antibodies and 0.25x10^6^ labeled cells/well were added to macrophages plate, placed in the Incucyte live-cell analysis system (Sartorius, Germany) and scanned every hour.

### Specificity profile

Membrane Proteome Array (MPA) screening was conducted at Integral Molecular, Inc. (Philadelphia, USA). The MPA is a protein library composed of 6,000 distinct human membrane protein clones, each overexpressed in live cells from expression plasmids. Each clone was individually transfected in separate wells of a 384-well plate followed by a 36 hours incubation (37). Cells expressing each individual MPA protein clone were arrayed in duplicate in a matrix format for high-throughput screening. Before screening on the MPA, the tested antibody concentration for screening was determined on cells expressing positive (membrane-tethered Protein A) and negative (mock-transfected) binding controls, followed by detection by flow cytometry using a fluorescently-labeled secondary antibody. The tested antibody was added to the MPA at the predetermined concentration, and binding across the protein library was measured on an Intellicyt iQue using a fluorescently-labeled secondary antibody. Each array plate contains both positive (Fc-binding) and negative (empty vector) controls to ensure plate-by-plate reproducibility.

### ImmunoGraft® Model

The study was conducted by Champions Oncology (New Jersey, USA). Four-week-old NOD/Shi-scid/IL-2Rγnull immunodeficient mouse strain (NCG, Charles River) mice were engrafted intravenously with cord blood-derived CD34^+^ hematopoietic stem and progenitor cells, two days after chemical myeloablative treatment (Busulphan). CD34^+^ cord blood cells (i.v. injection of 50x10^3^-100x10^3^ cells) from several donors were used and randomized across groups. Fourteen weeks after cell injection, engraftment level was monitored with the analysis of human CD45^+^ cells among total blood leukocytes (mouse and human) by flow cytometry. Only mice with a humanization rate (hCD45/Total CD45) above 25% were used in the study. Female human cord blood-derived CD34^+^ NCG mice implanted with human melanoma CTG-1769 tumors were allocated to two treatment groups of 12 mice each, as follows: G1 (IgG4 isotype control (25mg/kg)) and G2 (Anti-4CB1 (25mg/kg)). Each group consisted of 4 CBL (cord blood leukocytes) donors. hIgG4 isotype antibody (Sino Biological, #HG4K) served as a control. Antibodies were administered intravenously twice a week for a period of 4 weeks. During the treatment phase, animals were observed daily for clinical signs, body weight (twice a week), and tumor volume (twice a week) that was measured with digital caliper, and calculated using the formula: V = 0.5 a × b^2^ where a and b were the length and width of the tumor, respectively, and the TGI_TV_ (tumor growth inhibition by tumor volume) was calculated by the equation, TGI_TV_% =[1-(T_i_-T_0_)/(C_i_-C_0_)] × 100%. (T_i_: the mean tumor volume of the treated group at Day i following treatment; T_0_: the mean tumor volume of the treated group at Day 0; C_i_: the mean tumor volume of the control group at Day i following treatment; C_0_: the mean tumor volume of the control group at Day 0). Analysis was performed on data from day 13 post grouping (last day in which all mice were alive). At termination, the tumors of animals were weighed, average tumor weight in each group was determined, and the TGI_TW_ (tumor growth inhibition by tumor weight) was calculated by the formula: TGI_TW_% = (W_control group_ -W_treatment group_)/W_control group_×100%. W refers to the mean tumor weight.

### Humanized hBTLA and hBTLA/hHVEM mice models

The experiments were conducted by Biocytogen Pharmaceuticals (Beijing, China). The MC-38 murine colon carcinoma cell line was purchased from Shunran Shanghai Biological Technology Co., Ltd. The MC-38 cells were genetically modified to express human (h) HVEM in place of mouse (m) HVEM by Biocytogen Pharmaceuticals (Beijing) Co., Ltd. and named B-hHVEM MC-38 (Clone:2-A05). For the efficacy evaluation of Anti-4CB1 and its combination with Anti-mPD-1 in the treatment of Subcutaneous B-hHVEM MC38 Colon Carcinoma study, B-hBTLA mice; a C57BL/6 mice in which the exon 2 of mouse Btla gene was replaced by human BTLA exon 2, were subcutaneously inoculated with B-hHVEM MC38 tumor cells (1×10^6^) in 0.1 mL PBS in the right hind flank for tumor development. Seven days after tumor cell inoculation, animals with a mean tumor size of 103 mm^3^ were randomly enrolled into four study groups with ten mice per group. G1 (Isotype control, hIgG4 (10 mg/kg) + ratIgG2a (1 mg/kg)), G2 (Anti-4CB1 (10 mg/kg) + ratIgG2a (1 mg/kg)), G3 (hIgG4 (10 mg/kg) + Anti-mPD-1 (1 mg/kg)) and G4 (Anti-4CB1 (10 mg/kg) + Anti-mPD-1 (1 mg/kg)). Dosing started on the day of grouping (Day 0). All test articles were administered twice per week for a total of six times. The tumor volume and body weight were measured and recorded twice per week, using a caliper, and the volume was expressed in mm^3^ as described above. During the entire experiment, animals were monitored every day for their behavior and status. The study was terminated at day 53 post grouping. At the endpoint, TGI_TV_ was calculated and Kaplan-Meier survival curves were plotted.

For the evaluation of the efficacy of Anti-4CB1 in the treatment of the subcutaneous B-hHVEM MC38 colon carcinoma study, B-hBTLA/hHVEM mice; a C57BL/6 mice in which exon 2 of mouse Btla gene and exons 1∼6 of mouse Tnfrsf14 gene were replaced by the equivalent human exons, were subcutaneously injected with 5×10^5^ B-hHVEM MC-38 tumor cells per animal in the right front flank for tumor development. Seven days after tumor cell inoculation, animals with a mean tumor size of 90 mm^3^ were randomly enrolled into three study groups with ten mice per group; G1 (isotype control (25 mg/kg)), G2 (Anti-4CB1 (10 mg/kg)) and G3 (Anti-4CB1 (25 mg/kg)). All test articles were intravenously administered at a frequency of once every five days for a total of five times. The tumor volume and body weight were measured in two dimensions using a caliper as described above and recorded twice per week. The study was terminated at day 43 post grouping. At the endpoint, TGI_TV_ was calculated and Kaplan-Meier survival curves were plotted.

### Measurement of soluble HVEM in human serum

Measurement of human soluble HVEM (sHVEM) in human serum was performed using Human HVEM/TNFRSF14 DuoSet ELISA Kit (R&D Systems, #DY356), according to the manufacturer’s instructions. Briefly, plate was coated with Capture Antibody diluted in PBS for overnight at 4°C. Plate was washed and blocked with 1% BSA for one hour at room temperature followed by washes. Human serum and standard samples were diluted x4 with FBS (Gibco, #10270106) and incubated for two hours at room temperature. Plate was washed and incubated with Detection antibody for one hour at room temperature. Plate was washed and incubated for 30 minutes at room temperature with Streptavidin-HRP antibody, followed by washes and addition of TMB and stop solution. Absorbance was read at 450nM and 570nM.

### Immunohistochemistry of tumor micro-array (TMA) of responders and non-responders to Anti-PD1 treatment

Procedure was performed by Histospeck (Tel Aviv, Israel). 5 µm sections were prepared from each block of TMA, deparaffinized, and antigen retrieval was performed in a basic solution (pH=9). Triple staining was performed using a mouse monoclonal anti-CD3 antibody (Proteintech, #360181-1-Ig), a rabbit monoclonal anti-PD-L1 antibody (Cell Signaling, #13684) and a goat Human HVEM/TNFRSF14 polyclonal antibody (R&D Systems, #AF356) as primary antibodies. Anti-mouse Cy2, anti-rabbit Cy3 and anti-goat Cy5 secondary antibodies were used (Jackson; #715-545-151/152/147, respectively) and nuclei were stained with DAPI. Imaging was performed with a Leica TCS SP5 confocal laser-scanning microscope (X63 magnification) and image analysis was performed using the FIJI (Image J2) software.

### Statistics

All *in-vitro* experiments presented results are an average of at least 3 experiments. Data were analyzed using the paired or unpaired one or two-tailed Student’s t-test, as appropriate. *In-vivo* experiments results were represented by means and standard error of the mean (Mean ± SEM). Statistical analysis of TGI_TV_ and TGI_TW_ was performed using one-way ANOVA with Dunnett’s multiple comparisons test or using the one-tailed unpaired Student’s t-test. p ≤ 0.05 was regarded as statistically significant. For Kaplan-Meier survival curve Log-rank (Mantel-Cox) test were performed. In all graphs, error bars represent Standard Error. Asterisks indicate P values: *p ≤ 0.05, **p ≤ 0.01, ***p ≤ 0.001.

## Results

### Anti-4CB1 demonstrates specific binding to HVEM and activates T cells

In order to generate an anti-HVEM mAb that specifically blocks HVEM-BTLA interaction, we used phage display technique, as described in the Methods. Using flow cytometry and ELISA, we have identified scFv2 as a binder to HVEM and as a specific blocker for human HVEM-BTLA interaction (data not shown). Based on the binding and HVEM-BTLA blocking results, scFv2 was selected for conversion into full IgG by sub-cloning the variable region into IgG4 (S228P) backbone (named hereinafter Anti-4CB1). Anti-4CB1 underwent affinity maturation by targeting specific amino acids in the variable region sequence of the antibody.

Binding of Anti-4CB1 to recombinant hHVEM protein was re-confirmed by ELISA and EC_50_ was calculated using GraphPad Prism, as shown in Figure 1A. In addition, the ability of Anti-4CB1 to bind HVEM expressed on cells was re-confirmed by flow cytometry on CHO-K1 cells overexpressing the human, mouse and cynomolgus HVEM genes. CHO-K1 overexpressing HVEM cells were incubated with APC-labeled Anti-4CB1 or isotype control at different concentrations (5-100,000 ng/mL) for one hour on ice. Cells were washed and analyzed by flow cytometry. Effective concentrations (EC_50_) of Anti-4CB1 binding to HVEM protein expressed on CHO-K1 transfected cells was calculated using GraphPad Prism (Supplementary Figure S1A). The ability of Anti-4CB1 to bind endogenous HVEM was tested in IFNγ stimulated melanoma cells. Melanoma cells were first confirmed by staining with anti-melanoma (a cocktail of monoclonal antibodies against pmel17 and MART-1, data not shown), then, incubated with different concentrations (1-10,000 ng/mL) of APC labeled Anti-4CB1 or isotype control for one hour on ice. Expression was analyzed by flow cytometry and EC_50_ was calculated using GraphPad Prism (Supplementary Figure S1B).

**Figure 1.**
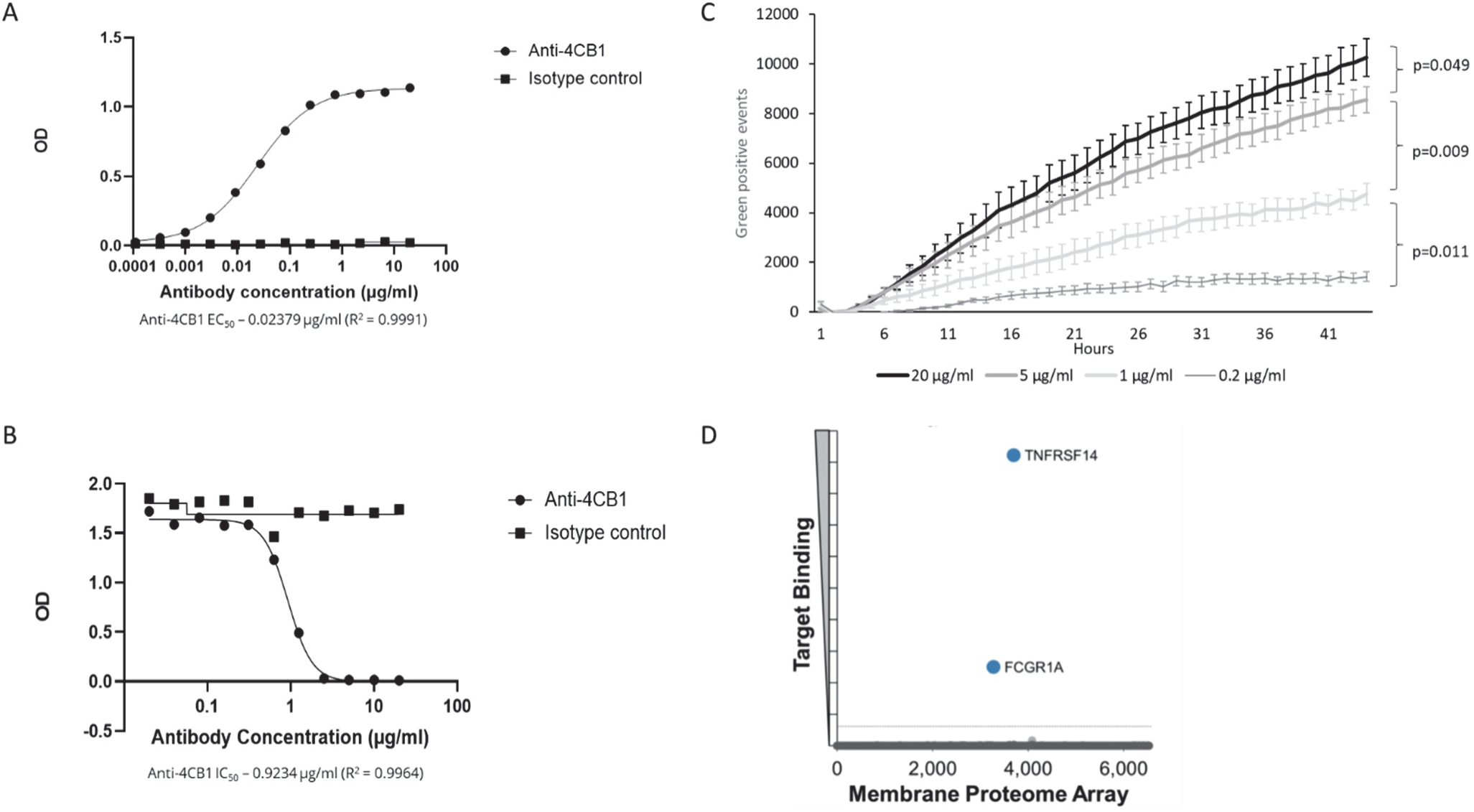
Anti-4CB1 demonstrates specific binding to HVEM and blocking of HVEM-BTLA interaction. (A) Binding of Anti-4CB1 or isotype control to human HVEM-His tag recombinant protein was tested in a range of concentrations 0.0001-20 μg/mL. The EC50 of 4CB1 was calculated using GraphPad Prism. (B) Blocking ELISA of human HVEM-BTLA interaction by Anti-4CB1 or isotype control tested at concentrations of 0.02-20 μg/mL. (C) CHO-K1 hHVEM/TCR activator cells were incubated with 0.2, 1, 5 and 20 μg/mL of Anti-4CB1 or isotype control, followed by addition of BTLA/NFAT eGFP Reporter Jurkat cells or medium. eGFP reporter activity was tested by Incucyte live-cell analysis system. Figure represents positive events with Anti-4CB1 treatment after subtracting isotype control positive events. (D) Membrane Proteome Array screen for specificity of Anti-4CB1. FCGR1A served as positive (FC-binding) control. Confirmed binding interactions are displayed in blue and any proteins that did not pass validation were removed.

To ensure that the blocking ability of Anti-4CB1 was not lost during affinity maturation, blocking ELISA was performed by incubating hHVEM with its main inhibitory receptor hBTLA in the presence of different concentrations (0.02-20 μg/mL) of Anti-4CB1 or isotype control (Figure 1B). Moreover, blocking of the hHVEM interaction with its additional interacting molecules, hLIGHT or hCD160, was assessed by ELISA. hHVEM was incubated with hLIGHT or hCD160 in the presence of different concentrations (0.02-20 μg/mL) of Anti-4CB1 or isotype control (Figures S2A and S2B, respectively). Results show that Anti-4CB1 completely abolished the major HVEM-BTLA interaction as compared to isotype control, as well as the interaction with the T-cell inhibitory receptor CD160, while no inhibition of the HVEM-LIGHT interaction was observed.

The ability of Anti-4CB1 to inhibit HVEM-BTLA interaction was also tested in cell-based co-inhibitory bioassay. First, we generated recombinant Jurkat T cells expressing eGFP gene under the control of NFAT response elements. Then, cells were transduced with CD272 (BTLA) for constitutive expression of human BTLA. In addition, we generated CHO-K1 cells overexpressing human HVEM with the TCR Activator Lentivirus. When these two cells are co-cultured, TCR complexes on effector cells are activated by the TCR activator on target cells, resulting in expression of the NFAT eGFP reporter. However, BTLA and HVEM ligation prevents TCR activation and suppresses the NFAT-responsive eGFP activity. Thus, BTLA or HVEM blocking antibodies reactivate the NFAT responsive eGFP reporter. To test the ability of Anti-4CB1 to block HVEM-BTLA interaction and activate the BTLA transduced Jurkat cells, CHO-K1 HVEM/TCR activator cells were incubated with Anti-4CB1 or isotype control antibodies at different concentrations (0.2-20 μg/mL), and co-cultured with BTLA/NFAT eGFP Reporter Jurkat cells for 45 hours. The results demonstrate a dose-dependent activation of the BTLA/NFAT eGFP Reporter Jurkat cells in the presence of Anti-4CB1 (Figure 1C).

Similarly, Jurkat T cells expressing eGFP gene under the control of NFAT response elements were transduced with CD160 for constitutive expression of human CD160. In addition, Dynabeads were coated with human HVEM protein. Activation of the CD160 transduced Jurkat cells was achieved by incubation with OKT3, resulting in expression of the NFAT eGFP reporter. However, ligation of CD160 with the HVEM-coated beads, suppresses the NFAT-responsive eGFP activity. As expected, the addition of 20 µg/mL Anti-4CB1, led to enhanced activation of the CD160/NFAT eGFP Reporter Jurkat cells, compared to isotype control (Supplementary Figure S2C).

To further evaluate the specificity of Anti-4CB1 and rule out any off-target antibody binding, which might lead to adverse events in clinical setup, we used the Membrane Proteome Array (Integral Molecular) screen. Results show that out of 6,000 human membrane proteins tested, Anti-4CB1 showed reactivity only against HVEM (TNFRSF14), suggesting high specificity of the antibody (Figure 1D). Overall, these results suggest that Anti-4CB1 is a specific anti-HVEM mAb, which specifically blocks the T-cell inhibitory HVEM-BTLA and HVEM-CD160 interactions.

### Anti-4CB1 enhances the killing ability of TILs towards autologous melanoma cells and increases the expression of activation markers CD137 (41BB) and CD107a

Since Anti-4CB1 binds HVEM and blocks its binding to BTLA, we hypothesized that it would alleviate the inhibitory signaling effect on T cells. Thus, specific killing of primary melanoma cells in the presence of autologous BTLA-expressing TILs was measured. Adhered melanoma cells were pre-incubated with Anti-4CB1 or isotype control and autologous TILs were pre-incubated with isotype control or Anti-PD1 antibody. Melanoma cells and TILs were then co-incubated in all possible combinations, and specific killing of melanoma cells was measured by detecting Caspase 3/7 events in time lapse microscopy. After 12 hours of co-culture, Anti-4CB1 alone and in combination with Anti-PD1 was found to increase total cell killing by 27% and 64%, respectively, as compared to cells treated with an isotype control (Figure 2A). These results demonstrate that Anti-4CB1 has the ability to enhance the cytotoxicity of potent TILs towards melanoma cells, while combination with Anti-PD1 can further enhance the cytotoxicity effect.

**Figure 2.**
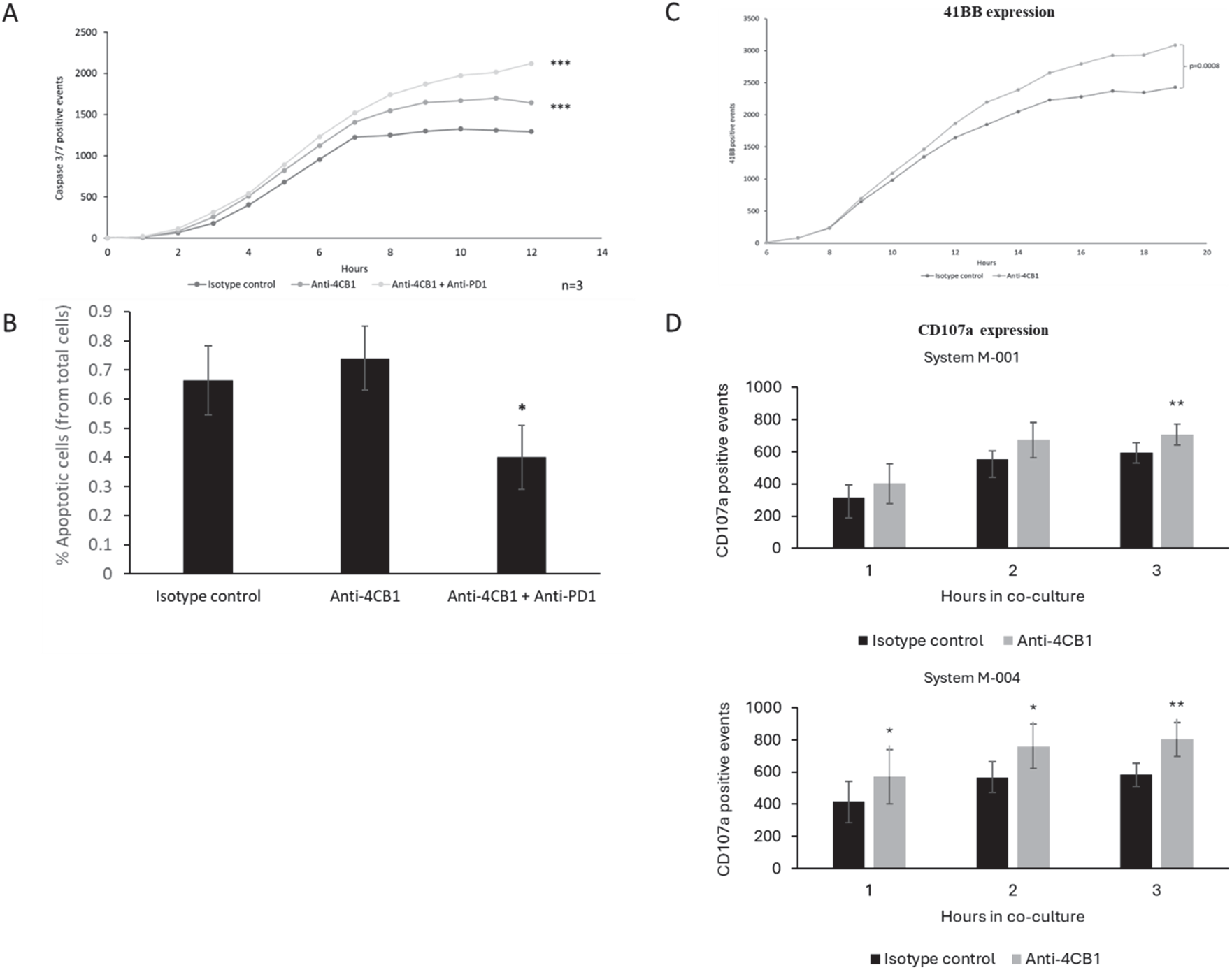
Anti-4CB1 enhances T-cell activation. (A) Killing assay with Anti-4CB1 in combination with Anti-PD1. Melanoma cells were pre-incubated with 20 µg/mL Anti-4CB1 or isotype control followed by co-culture with autologous TILs pre-incubated with of Anti-PD1 or isotype control. Killing of melanoma cells by TILs, reflected by positive Caspase 3/7, was measured in time lapse microscopy. (B) Apoptosis of melanoma cells. Melanoma cells were pre-incubated with 20 µg/mL Anti-4CB1, isotype control or combination of Anti-4CB1 and 20 µg/mL Anti-PD1. Positive Caspase 3/7 was measured after 12 hours. (C-D) Expression of TILs activation markers. TILs were incubated with 20 µg/mL Anti-4CB1 antibody or isotype control treated melanoma cells, then (C) labeled 41BB antibody was measured for 19 hours in time lapse microscopy, (D) or conjugated CD107a antibody for 3 hours.

Ruling out the possibility that the increased cytotoxicity was directly caused by the antibody, and not due to increased T cell killing, the direct cytotoxic effect of Anti-4CB1 alone, or in combination with Anti-PD1, on melanoma cells was measured. Very low percentages of apoptotic cells were observed. Most importantly, no increase in apoptotic cells was observed in melanoma cells that were incubated with Anti-4CB1, alone, or in combination with Anti-PD1 (Figure 2B), indicating that the antibody itself is not cytotoxic and that the increased cancer killing observed was rather due to increased T cell cytotoxicity.

We next decided to test whether the enhanced cytotoxicity, observed following the addition of Anti-4CB1 to melanoma-TIL co-culture, is indeed mediated by the T-cells. To this end, we examined the expression of 41BB and CD107a, as T-cell activation markers. TILs were incubated with Anti-4CB1 or isotype control pre-treated melanoma cells and the expression of 41BB was monitored over 19 hours of co-culture. Figure 2C shows that the number of activated TILs was increased by 27% in the presence of Anti-4CB1, compared to isotype control suggesting that the enhanced cytotoxicity is a result of TILs activation. Similarly, the expression of the activation marker CD107a was measured in these cells and Anti-4CB1 led to 19% (p=0.01) increase in CD107a positive cells after 3 hours of co-culture, as compared to isotype control. When a second melanoma cell system was tested, the same trend was observed in the presence of Anti-4CB1 – a 37% (p=0.01) increase in CD107a positive cells after 3 hours of co-culture, as compared to isotype control, suggesting, once again an effect on TILs activation (Figure 2D).

Having shown that the cytotoxicity of TILs can be enhanced using Anti-4CB1, the ability to enhance macrophages was also tested. Monocytes were isolated from healthy donors and matured into M1 macrophages. Expression of macrophages markers was confirmed by flow cytometry (Supplementary Figure S3A). HVEM-expressing A549 lung cells and M1 macrophages were co-incubated with Anti-4CB1, Anti-PD1, their combination or with isotype control. Specific phagocytosis of A549 cells in the presence of M1 macrophages was measured by detecting pHrodo red dyed events in time lapse microscopy. After 14 hours of co-culture, Anti-4CB1 or Anti-PD1 alone were found to increase total cell phagocytosis by 54% and 84%, respectively, without reaching statistical significance, as compared to cells treated with isotype control. However, Anti-4CB1 in combination with Anti-PD1 was found to increase total cell phagocytosis by 300% (p=0.04), as compared to isotype control (Supplementary Figure S3B).

Overall, these results indicate that binding of Anti-4CB1 to HVEM expressed on cancer cells leads to increased cytotoxicity, as a result of increased TILs activation. In addition, the combination of Anti-4CB1 and Anti-PD1 increased macrophage-mediated phagocytosis.

### Anti-4CB1 enhances cytotoxicity *ex-vivo* in a wide range of malignancies including cases in which Anti-PD1 does not

The efficacy of Anti-4CB1 was further evaluated in a series of 49 ex-vivo cancer samples derived from freshly digested tumor tissues, which include both tumor cells and cells from their microenvironment. Supplementary Table S1 provides the sample numbers and corresponding indications. Immediately after tissue digestion, cells were co-cultured with autologous PBMCs in the presence of Anti-4CB1, Anti-PD1 or isotype control. Cell death of cancer cells was measured by Caspase-3/7 positive events. Figure 3A depicts the percentage change in total cell killing following incubation with Anti-4CB1 in comparison to the isotype control. Enhanced killing, defined as a minimum of 30% change in total cell killing and p-value ≤ 0.05, was observed in 14 out of 49 samples (28.5%). In contrast. incubation with Anti-PD1 resulted in increased killing in only 7 out of 49 samples (14.3%), as illustrated in Figure 3B. Notably, among the samples exhibiting enhanced killing, only 5 samples showed enhanced killing to both Anti-4CB1 and Anti-PD1, meaning that 9 samples were unique to Anti-4CB1 while only 2 samples were unique to Anti-PD1. Analyzing the results by indication revealed higher killing rates following treatment with Anti-4CB1 compared to treatment with Anti-PD1 was observed in colon cancer (44% vs. 11%), melanoma (75% vs. 25%), ovarian cancer (20% vs. 0%) and renal cancer (18% vs. 8%) (Supplementary Table S2).

**Figure 3.**
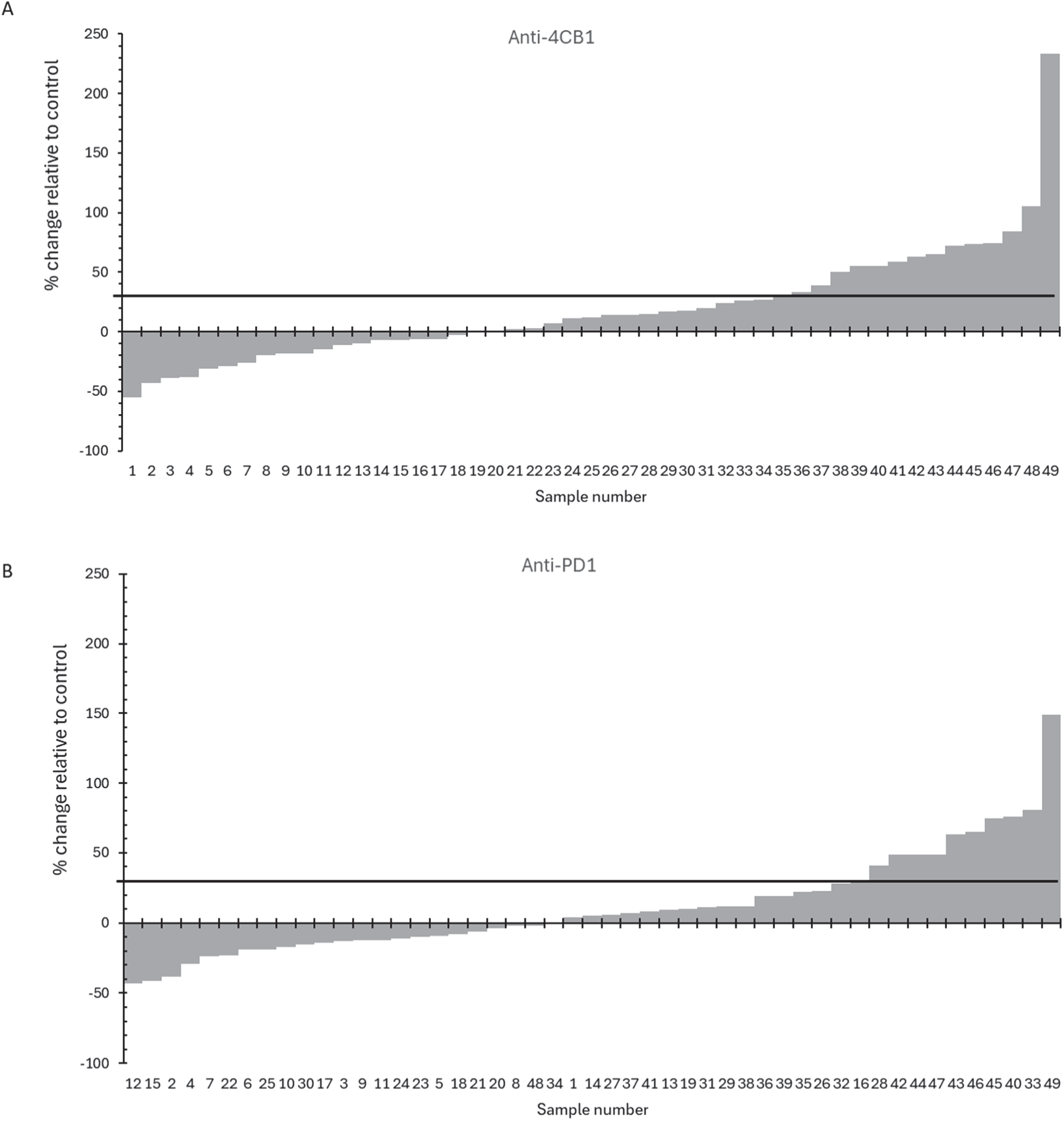
Anti-4CB1 enhances cytotoxicity in a variety of cancer tissues *ex-vivo*. Freshly isolated cells from bladder cancer (n=2), colon cancer (n=9), endometrial cancer (n=7), gastric cancer (n=1), hepatocellular carcinoma (n=5), lung cancer (n=3), melanoma (n=4), ovarian cancer (n=5), pancreatic cancer (n=2) and renal cancer (n=11) tissues were co-cultured with autologous PBMCs in the presence of (A) 5-20 µg/mL (depending on the tissue) of Anti-4CB1, (B) 20 µg/mL Anti-PD1 or isotype control (with the equivalent concentration). Killing of cancer cells, reflected by positive Caspase 3/7, was measured in time lapse microscopy. Specific killing was calculated by subtracting the positive events in PBMCs only wells from the positive events in cancer cells with PBMC wells. Figures show percent change in total cell killing relative to treatment with isotype control.

All together these results suggest that Anti-4CB1 has an effect on cytotoxicity in a wide range of cancer tissues. Moreover, Anti-4CB1 as monotherapy, may be efficacious in cases where Anti-PD1 is not, underscoring the potential of Anti-4CB1 as a promising therapeutic approach.

### Anti-4CB1 reduces tumor growth in colon and melanoma cancer *in-vivo* alone and in combination with Anti-PD1

In light of the cytotoxicity results *in-vitro*, we wanted to test the function and efficacy of Anti-4CB1 alone and in combination with a murine Anti-PD1 *in-vivo*. We have shown that Anti-4CB1 binds to CHO-K1 cells overexpressing mouse HVEM (Supplementary Figure S1A) and confirmed this binding also to recombinant mouse HVEM by ELISA (Supplementary Figure S4A). Having shown that Anti-4CB1 blocks the human HVEM-BTLA interaction (Figure 1B), we wanted to test whether it also blocks the murine HVEM-BTLA interaction. mHVEM was incubated with mBTLA in the presence of Anti-4CB1 or isotype control, however results show that although Anti-4CB1 binds murine HVEM it does not disrupt the murine HVEM-BTLA interaction over a range of antibody concentrations (Supplementary Figure S4B). Hence, we decided to first test the efficacy of the antibody in the treatment of subcutaneous B-hHVEM MC-38, in transgenic B-hBTLA mice (expressing human BTLA instead of the murine BTLA under the same endogenous promoter). Four groups with ten mice per group were tested, as detailed in the Methods. On day 21 post grouping, the mean tumor volume of isotype control group was 1236±193 mm^3^, and that of the Anti-4CB1, Anti-mPD1 and the combination groups was 979±189 mm^3^, 483±234 mm^3^ and 148±36 mm^3^ respectively. The corresponding TGITV were 23% (not significant), 66% (p=0.0115), and 95% (p<0.0001) (Figure 4A and Supplementary Table S3A) as compared to isotype control. The median survival days of groups G1-G3 were 28, 30 and 43, respectively. Most importantly, the median survival time for the combination of Anti-4CB1 with Anti-mPD1 was not reached and it significantly prolonged the survival time of animals compared with the Anti-mPD1 group alone (p=0.0175) (Figure 4B). Given the observed anti-tumor activity also in the Anti-4CB1 group, we aimed to evaluate the *in-vivo* efficacy of Anti-4CB1 alone at two different concentrations in treating subcutaneous B-hHVEM MC38 colon carcinoma in a transgenic B-hBTLA/hHVEM mice model. Three groups with ten mice per group were tested: Isotype control (25 mg/kg), Anti-4CB1 (10 mg/kg) and Anti-4CB1 (25 mg/kg). On day 25 post grouping, the mean tumor volume of G1-G3 were 1572±188 mm^3^, 1315±176 mm^3^ and 832±94 mm^3^, respectively, with TGI_TV_ of 17% (not significant) and 50% (p=0.0096), respectively, as compared to isotype control (Figure 4C, and Supplementary Table S3B). Kaplan Meier curve shows that Anti-4CB1 (25 mg/kg) treatment prolonged survival significantly compared to mice in the control group (p < 0.01), and the median survival days of G1-G3 were 32, 32 and 39 with 0%, 10% and 30% survival, respectively (Figure 4D).

**Figure 4.**
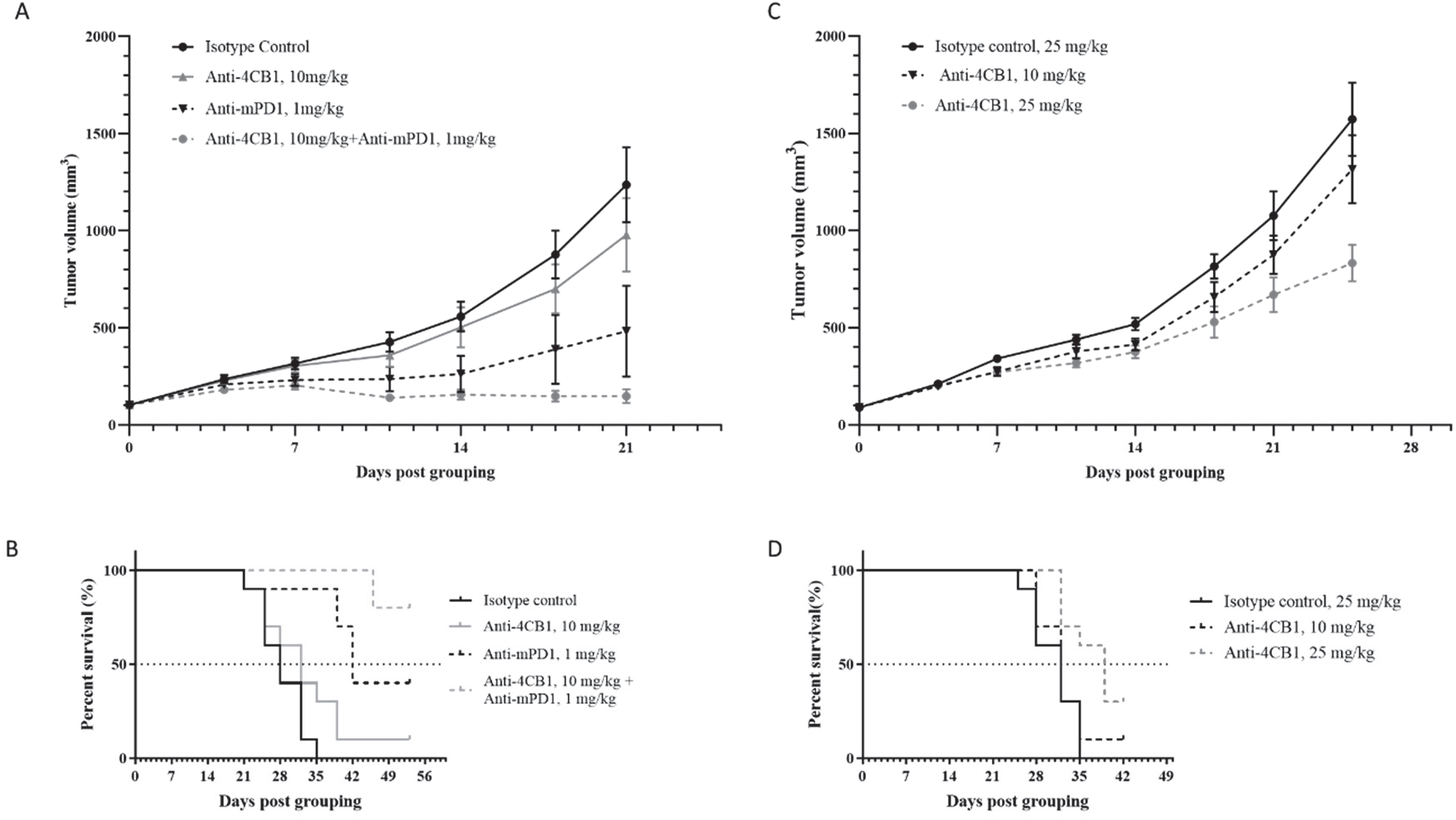
Anti-tumor response upon treatment with Anti-4CB1 or Anti-PD1 in transgenic mice model. (A) Mean tumor volume (mm^3^) upon treatment in B-hBTLA mice inoculated with B-hHVEM MC-38 cells. (B) Kaplan–Meier survival curve in B-hBTLA mice model. (C) Mean Tumor volume (mm^3^) upon treatment in B-hBTLA/hHVEM mice inoculated with B-hHVEM MC-38 cells. (D) Kaplan–Meier survival curve in B-hBTLA/hHVEM mice model.

In addition, the efficacy of Anti-4CB1, as a monotherapy was evaluated in a PDX humanized mouse model. Human cord blood-derived CD34^+^ boosted NCG mice were implanted with human melanoma (CTG-1769) tumors. When the mean tumor size reached 85 mm^3^, mice were divided into two treatment groups, as described in the Methods. HVEM expression on CTG-1769 melanoma cells was previously tested and the expression was at average of 61.45%. (data not shown). The experiment was terminated on day 23 post grouping. Since we observed that the response is donor dependent, the mean tumor volume of each CBL donor was calculated and compared between treatment groups. One CBL donor showed statistically significant anti-tumor response. On day 13 post grouping, the mean tumor volume of G1 was 915±73 mm^3^ and the mean tumor volume of G2 was 465±67 mm^3^, with a TGI_TV_ of 53% (p=0.0102). The corresponding TGI_TW_ was 35% (p=0.005), compared to isotype control treatment (Figure 5A-B and Supplementary Table S3C). No effect was observed in the other CBL donors (data not shown).

**Figure 5.**
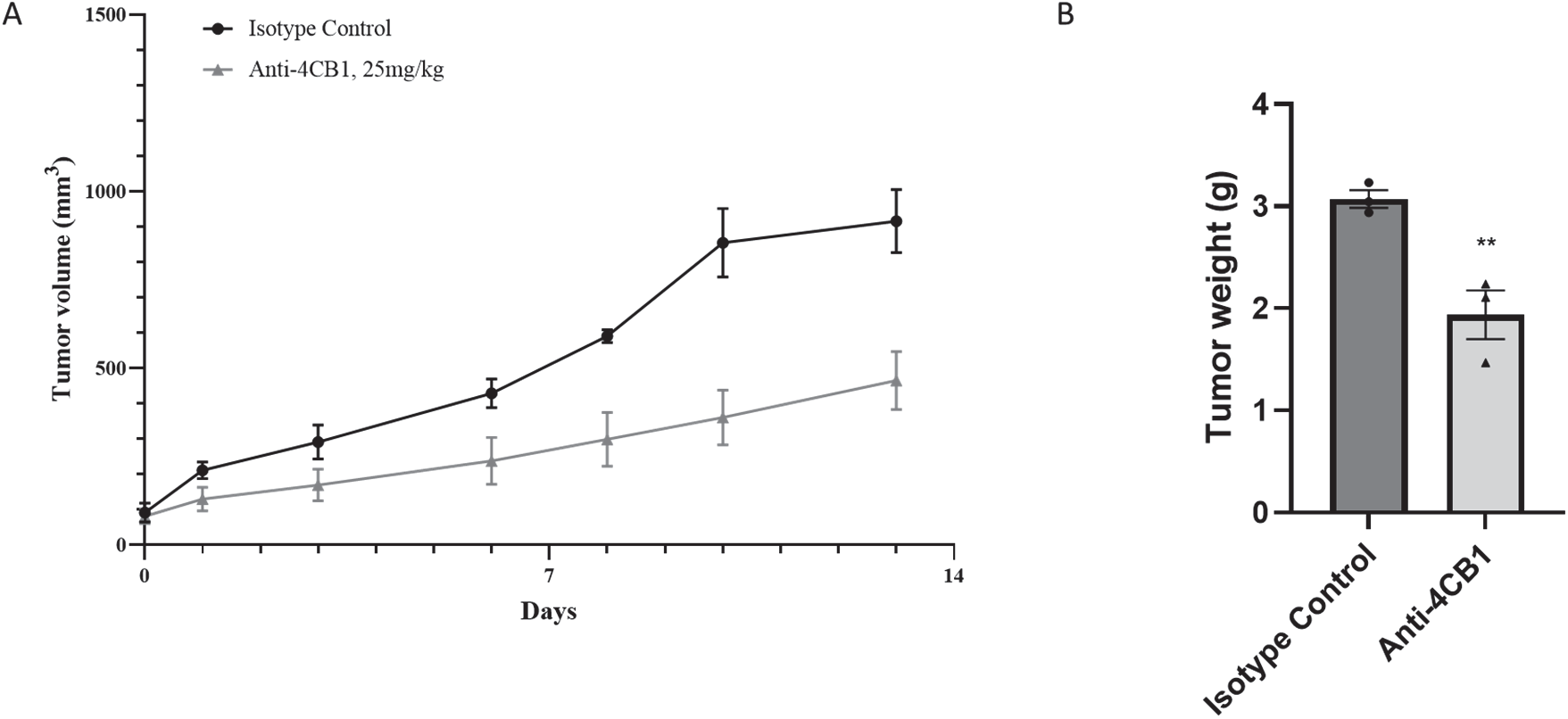
Tumor volume and tumor weight upon treatment with Anti-4CB1 in humanized mice model. (A) Mean tumor volume (mm^3^) upon treatment in myeloid-boosted humanized NCG mice inoculated with CTG-1769 melanoma model. (B) Mean tumor weight (g) on the day of sacrifice.

Overall, these experiments demonstrate the potential of Anti-4CB1 mAb as an effective monotherapy, outperforming the isotype control. Furthermore, when used in conjunction with Anti-PD1, enhanced anti-tumor activity was observed, surpassing that of Anti-PD1 alone.

### HVEM expression may serve as a marker for response to anti-HVEM and anti-PD1 immunotherapy

Finally, to investigate the correlation between HVEM expression and the response to ICI treatments, two distinct approaches were employed. First, sHVEM levels were quantified in serum samples obtained from both healthy donors and cancer patients. The serum samples from cancer patients were collected (when available) from individuals whose tumor tissues were tested in ex-vivo experiments. The results indicated significantly higher levels of sHVEM in cancer patients compared to healthy donors (p=0.0374), as shown in Figure 6A. Furthermore, when analyzing sHVEM levels in serum samples solely from cancer patients, it was observed that samples who did not respond to Anti-4CB1 treatment in our ex-vivo experiments (as defined above) exhibited significantly higher sHVEM levels (p=0.0463) compared to those who did respond (Figure 6B).

**Figure 6.**
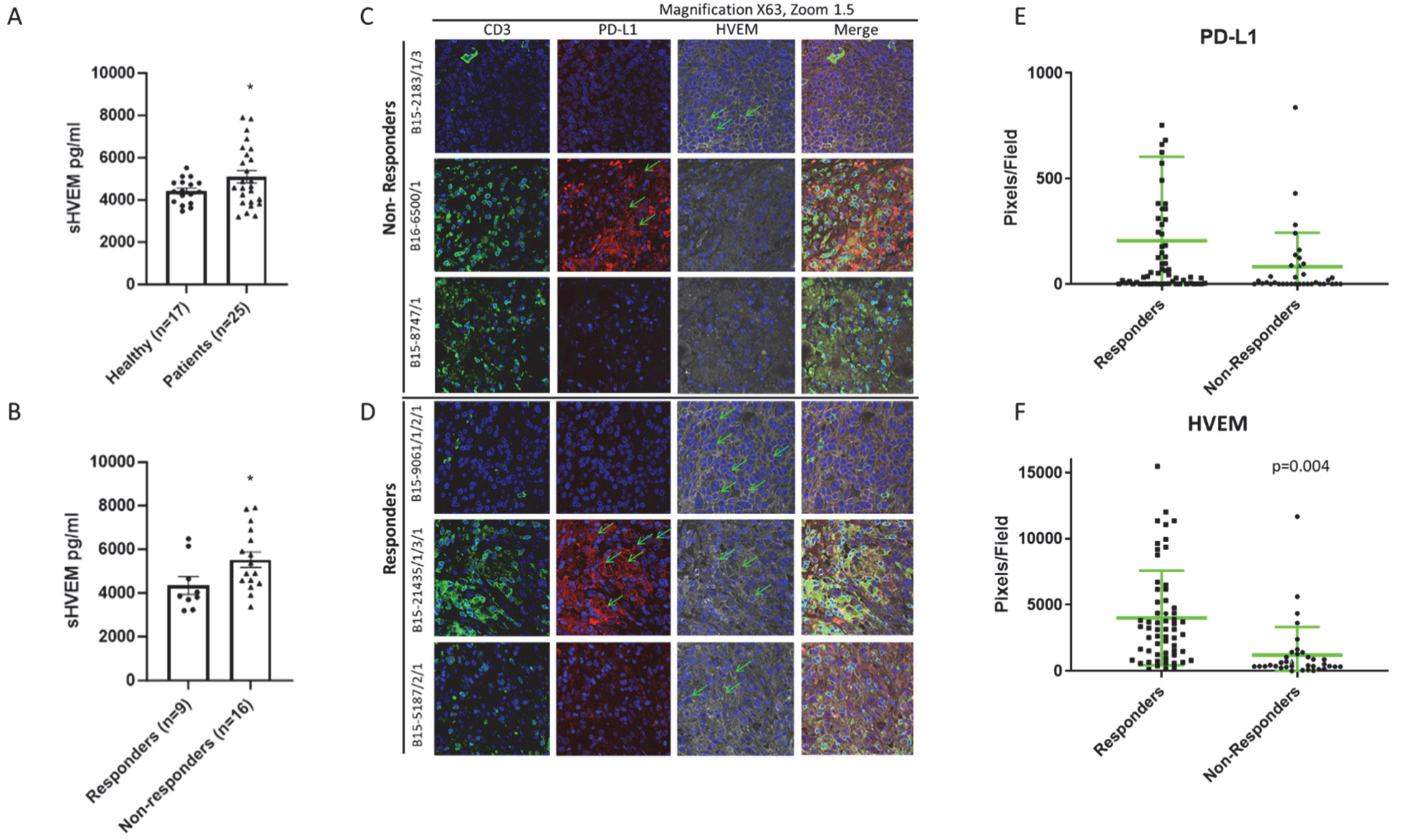
HVEM expression may serve as a marker for response to HVEM and PD1 immunotherapy. (A) Levels of soluble HVEM, measured by ELISA, in serum samples from healthy donors and cancer patients. (B) Levels of soluble HVEM, measured by ELISA, in serum samples from cancer patients in samples responding and non-responding to Anti-4CB1 treatment as defined in ex-vivo experiments. (C-D) IHC staining of CD3 (green), PD-L1 (red) and HVEM (grey) of melanoma tissues from patients who (C) did not respond to Anti-PD1 treatments and (D) patients who responded to Anti-PD1 treatment. Nuclei were stained with DAPI. (E) PD-L1 and (F) HVEM expression per sample, in responders and non-responders to Anti-PD1 treatment, as measured by total positive pixels/field.

Secondly, TMAs of melanoma tumor tissues were IHC stained for CD3, PD-L1 and HVEM expression. The TMAs contained 20 patients clinically characterized as responders and 16 patients clinically characterized as non-responders to PD1 blockade (Nivolumab, BMS or Pembrolizumab, Merck). The TMAs contained 1-3 cores from each of these patients. Representative pictures of CD3, PD-L1, HVEM and an overlay of the three staining are shown in Figure 6C-D. Staining of non-responders found a high degree of heterogeneity among the samples, with CD3, PD-L1 and HVEM all ranging from negative in some samples to highly expressed in others (Figure 6C). The same heterogeneity was observed in the responders’ samples (Figure 6D). However, quantification of the expression of each protein over the entire population of responders versus non-responders yielded surprising results. CD3 expression was comparable between the two populations and a small difference observed was not statistically significant (data not shown). PD-L1 expression was elevated in the responders’ population however this increase was not significant (Figure 6E). Remarkably, statistically significant (p=0.004) higher levels of HVEM were observed in the responders’ group as compared to the non-responders (Figure 6F).

These results suggest that sHVEM expression may distinguish between responders and non-responders to Anti-HVEM treatment. Additionally, tumor HVEM expression discriminates more effectively between responders and non-responders to Anti-PD1 treatment compared to PD-L1 expression. Therefore, HVEM expression may potentially serve as a valuable predictive marker for responses to ICI-based immunotherapies.

## Discussion

Recent years have seen significant advancements in cancer treatment with the development of drugs targeting immune checkpoint molecules. Despite these successes, only a limited proportion of patients benefit from these therapies, and the emergence of acquired resistance is relatively common. Therefore, there is an urgent need to develop new immune checkpoint agents to overcome these limitations.

In this study, we describe the development of a novel fully human monoclonal antibody that targets HVEM, a known ligand of the immune checkpoint BTLA. Given HVEM’s dual functional activity, both as a co-stimulatory and as an inhibitory molecule, targeting HVEM with a specific antibody could potentially induce an anti-tumor effect through various mechanisms. Our findings reveal that our proprietary Anti-4CB1 mAb can completely abolish the interaction between HVEM and both BTLA and CD160, without interfering with the interaction between HVEM and LIGHT, suggesting that the stimulatory function of HVEM should not be affected by the antibody.

Our *in-vitro* functional assays, together with the ELISA results, support the assumption that the cytotoxic ability of T-cells is enhanced by alleviating the inhibitory signal of BTLA and CD160, which is achieved by blocking the HVEM-BTLA and HVEM-CD160 interactions with Anti-4CB1. While it could be that the enhanced Caspase 3/7 expression observed in the killing assays is due to a direct effect of Anti-4CB1 on melanoma cells, we have shown that binding of the mAb to the melanoma cells does not induce apoptosis *in-vitro.* Although there is a difference between the activity of a blocking antibody and downregulation by RNAi, others have also shown that silencing of HVEM did not induce apoptosis of cancer cells (38,39). The assumption that Anti-4CB1 induces T-cell mediated immune response was further reinforced by the observed increased expression of 41BB and CD107a, as T-cell activation markers, after co-culture with melanoma cells. The effect of HVEM expressed in cancer cells on T-cell activation was also demonstrated by Zhang *et al*., who showed that co-culture of activated T-cells with HVEM-silenced ovarian cancer cells led to increased cytokine secretion (39). Aubert *et al*., showed that treatment with a murine anti-hHVEM mAb decreased tumor growth of prostate cancer and melanoma in humanized NSG mice (34). In addition to the T-cell mediated immune response, our study demonstrates that Anti-4CB1, when combined with Anti-PD1, enhances macrophage-mediated phagocytosis. This finding suggests that it may also impact the innate immune system.

The anti-tumor activity of Anti-4CB1 was further confirmed in a variety of *ex-vivo* experiments, suggesting its potential clinical effectiveness across multiple cancer indications. Most importantly, Anti-4CB1 demonstrated efficacy in twice as more patient samples compared to most promising immune-checkpoint inhibitor, Anti-PD1, including in cases where Anti-PD1 was ineffective. These findings are particularly noteworthy, as they indicate that Anti-4CB1 may benefit patient subpopulations that do not respond to existing therapies.

In line with our *ex-vivo* findings, tumor growth inhibition with Anti-4CB1 alone, was observed in our colon cancer transgenic model and in humanized mice using melanoma PDXs, suggesting once again the clinical relevance of the antibody. Similar results were previously shown with MC-38 colorectal cancer cells using a murine anti-HVEM antibody (33). Of importance, unlike Anti-4CB1, a surrogate antibody of Relatlimab, an anti-LAG3-blocking mAb, which has been recently approved for treating melanoma in combination with PD1-blocking Nivolumab, showed no anti-tumor efficacy as monotherapy in a MC38 colon carcinoma model (40).

An unprecedented number of clinical trials are evaluating the clinical efficacy of Anti-PD1 treatment in combination with other agents (41). Thus, we thought to test the effect of Anti-4CB1 in combination with Anti-PD1, the leading treatment for advanced melanoma and other cancers. *In-vivo* studies show superior anti-tumor activity of Anti-PD1 and Anti-4CB1 combination compared to either antibody alone. These findings are consistent with *in-vitro* results shown by Demerle *et al*., demonstrating that a murine anti-HVEM antibody worked synergistically with anti-PD-L1 to enhance the activation of T-cells in a lung cancer cell line (33). It has been described that compared to PD1, BTLA recruits additional phosphatases to inhibit T-cell signaling; while both PD1 and BTLA employ SHP2 to dephosphorylate CD28, BTLA employs also SHP1 to dephosphorylate both TCR and CD28 (42). Given a different mechanism of action, Anti-HVEM treatment may have an advantage in cases where Anti-PD1 has no clinical benefit or could have an improved effect when combined with Anti-PD1. Indeed, our work demonstrates the possibility of such enhanced effects. Moreover, blocking HVEM might have an advantage over antibodies blocking BTLA, due to the fact that HVEM interacts with additional immune checkpoints (such as CD160). This additional blockade, preventing multiple inhibitory signals that suppress T-cell activity, as demonstrated by Anti-4CB1, could improve efficacy and potentially address resistance issues seen in single-target therapies.

Since only 20-60% of patients show clinical benefit from ICIs treatments, there is constant search after predictive biomarkers to optimize patient selection. One of the many explored markers for Anti-PD1/PD-L1 response is the expression of PD-L1 in tumor cells. However, few challenges still exist including low expression, preventing PD-L1 from being a comprehensive and independent biomarker. Thus, identifying new predictive biomarkers is essential (43–45). It has been suggested that HVEM is more widely expressed than PD-L1 in melanoma cells and that high expression of HVEM is a marker for poor prognosis (46). In addition, Aldahlawi *et al*. showed that patients with malignant breast cancer exhibited a significant increase in both genetic and serum HVEM levels compared with non-malignant control subjects, suggesting its potential as biomarker (47). Others have suggested that circulating HVEM levels do not correlate with immunotherapy response (48) while in a recent review, it has been suggested that levels of the soluble immune checkpoints sTIM3, sBTLA4, sHVEM, sCTLA4, sPD-L1 and sPD-1 were higher after treatment with Nivolumab in non-responding versus responding patients with NSCLC (49). Interestingly, in our study, IHC staining of a TMA consisting of melanoma samples from patients treated with Anti-PD1 showed that HVEM expression could predict response to Anti-PD1 even better than PD-L1 expression. These results are in agreement with a previous study showing an association between high RNAseq HVEM expression and a favorable response to immune check blockade (50). As anticipated, our findings demonstrated higher sHVEM expression in cancer patients compared to healthy donors, consistent with previous studies (49). Remarkably, our results revealed significantly elevated sHVEM levels in serum samples from patients whose samples did not respond to our Anti-HVEM treatment *ex-vivo* compared to those who did respond. To the best of our knowledge, no studies have examined the relationship between baseline sHVEM levels and clinical response to HVEM blocking or other ICIs. Despite the limitation of a small sample size in the present study, if these findings are ratified in a larger cohort of patients, HVEM could be considered as another potential biomarker for predicting responses to Anti-PD1 and, more importantly, Anti-HVEM treatment.

To the best of our knowledge, to date, no HVEM blocking antibodies have reached the clinical stage. Anti-4CB1’s specificity to HVEM, combined with its blocking activity and efficacy in inducing an immune response, positions it as a promising candidate for development as an anti-cancer therapeutic agent. Its unique characteristics and potential to address current treatment limitations underscore the importance of further clinical investigation.

## Acknowledgements

The authors would like to thank the Sheba Medical Center Biobank and the Rabin Medical Center Biobank teams for their administrative support. The authors would like to thank Dr. Itay Spektor for assisting with the histological work.

## Ethics

All studies involving patient-derived material were approved by the Institutional Review Board of Sheba Medical Center and Rabin Medical Center, Israel.

## Author contributions

G.G.H and E.M.S conceived and designed the work and drafted the manuscript; M.S performed the experiments and assisted in manuscript writing; R.B, N.D, S.S.O and K.S performed the experiments and data analysis; A.L.B, G.H.S, J.S, E.S, R.E, I.B, S.G, Y.F, E,Y, S.E, assisted with data and specimen collection; G.M and E.G conceived the work and revised the manuscript.

