## Supplementary figures and tables for "Novel immunotherapy for multiple solid cancers using an Anti-HVEM blocking monoclonal antibody"

**Supplementary figures and legends**

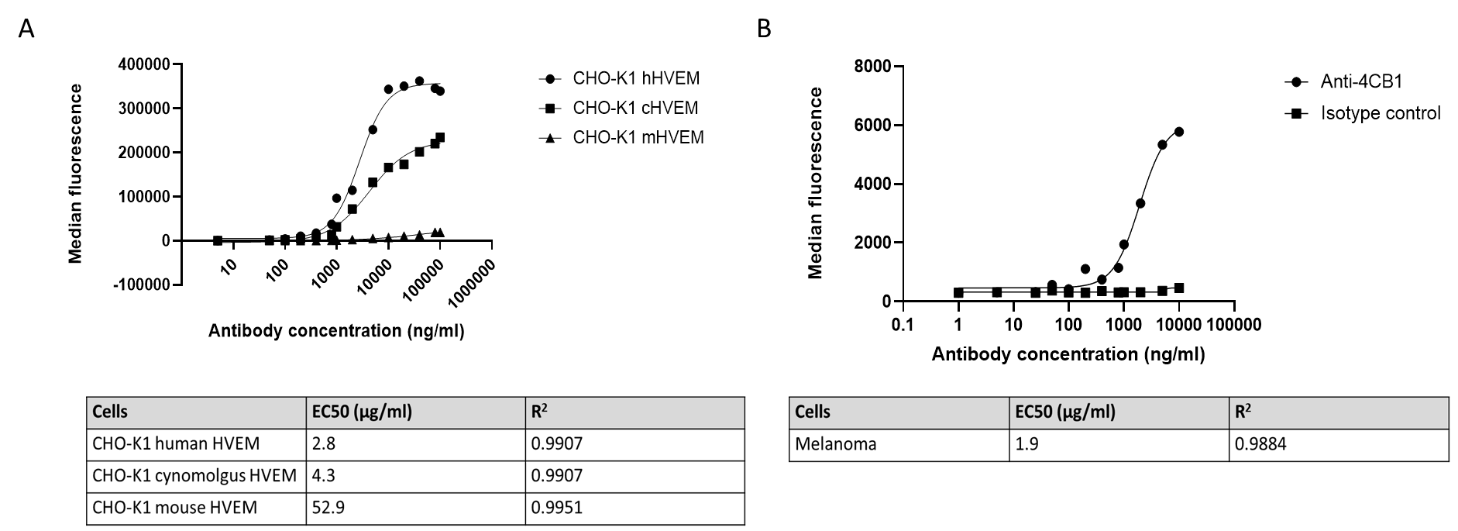

**Figure S1. Anti-4CB1 binds to surface HVEM expressed on cells.** Binding of APC labeled Anti-4CB1 to (A) CHO-K1 cells (5-100,000 ng/mL) expressing human, mouse or cynomolgus HVEM and (B) IFNγ stimulated melanoma cells (1-10,000 ng/mL).

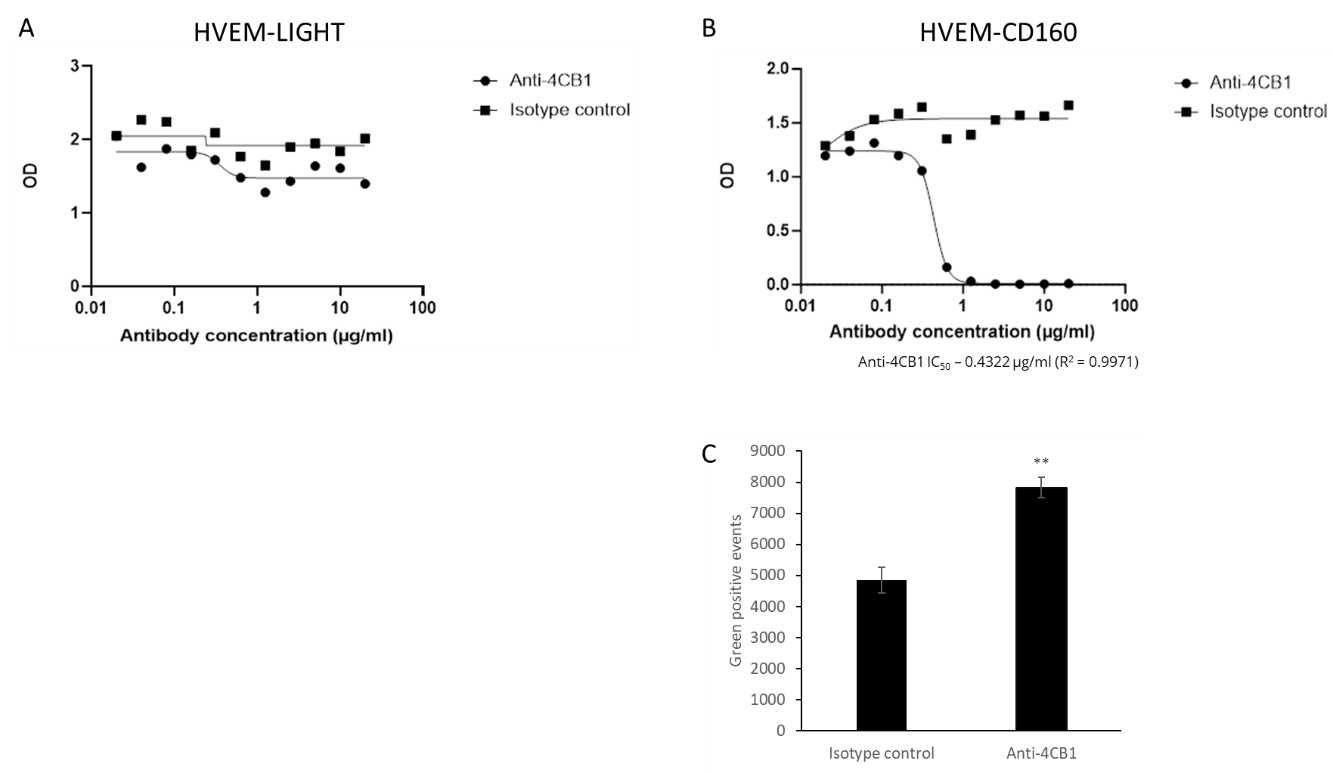

**Figure S2. Anti-4CB1 blocks human HVEM-CD160 interaction but not human HVEM-LIGHT interaction.** Blocking ELISA of (A) human HVEM-LIGHT and (B) human HVEM-CD160 interaction by Anti-4CB1 or isotype control tested at concentrations of 0.02-20 μg/mL. (C) hHVEM coated Dynabeads were pre-incubated with 20 μg/mL of Anti-4CB1 or isotype control followed by incubation with OKT3 stimulated Jurkat CD160/NFAT-eGFP cells. eGFP reporter activity was tested by Incucyte live-cell analysis system.

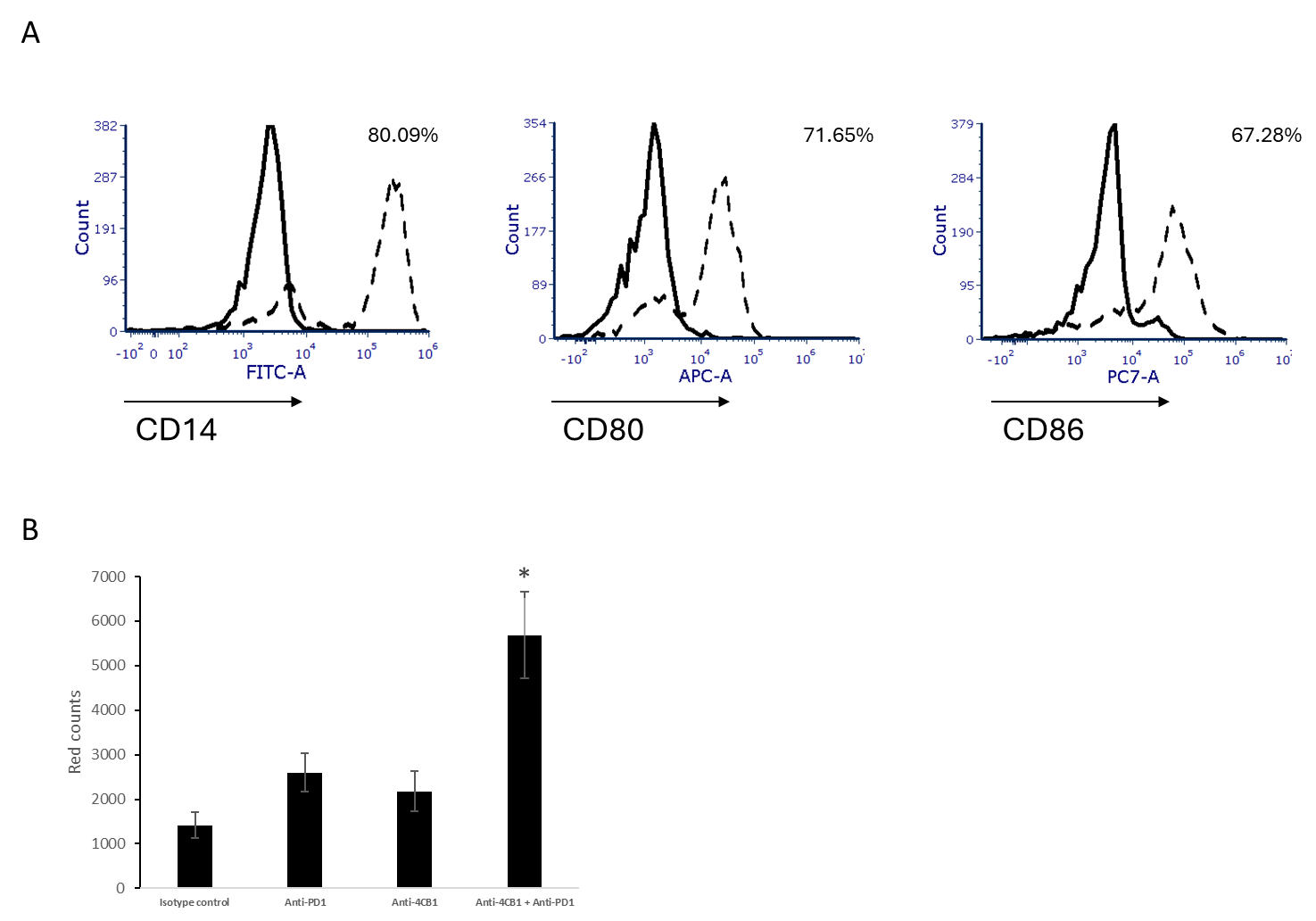

**Figure S3. Combination of Anti-4CB1 and Anti-PD1 increases phagocytosis**. (A) Flow cytometry staining of macrophages for the expression of CD14, CD80, CD86. (B) Macrophages and pHrodo Red cell labeled A549 cells were co-cultured with 20 µg/mL isotype control or Anti-4CB1 with or without Anti-PD1. Red fluorescence was measured to detect phagocytosis events in time lapse microscopy.

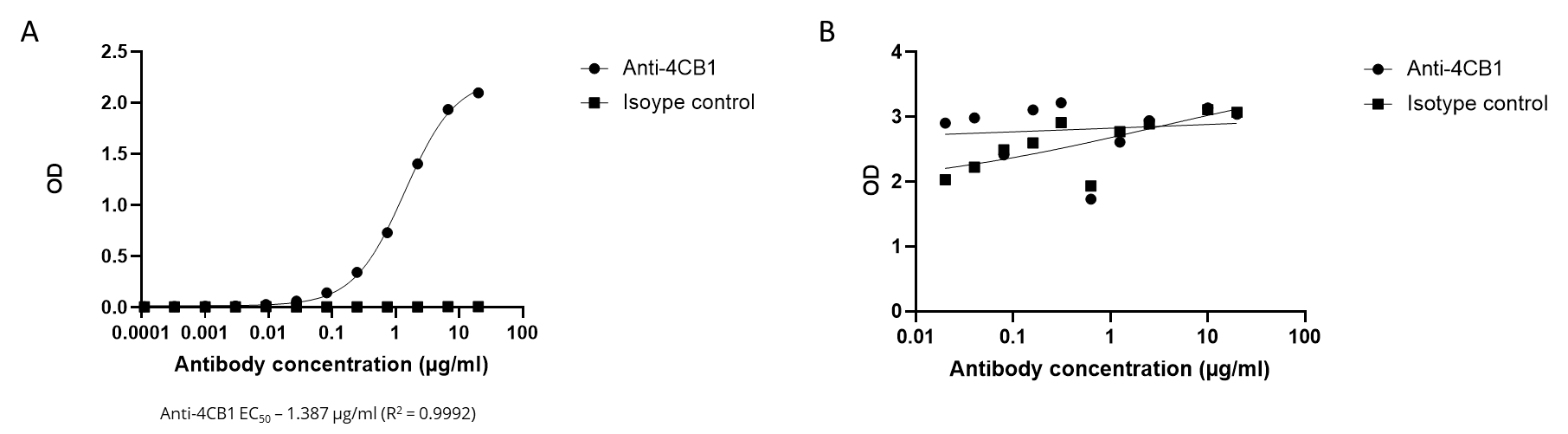

**Figure S4. Binding and blocking of Anti-4CB1 to murine HVEM** (A) Binding: Plate was coated with 1 μg/mL of mouse HVEM-His tag Protein or PBS for overnight at 4°C, then incubated with Anti-4CB1 mAb or isotype control in concentrations of 0.0001-20 μg/mL for one hour followed by incubation with peroxidase anti-human FC for 30 minutes and addition of TMB and stop solution. Absorbance was read at 450nM and 570nM. Effective concentrations (EC50) of Anti-4CB1 mAb binding to mouse HVEM-His tag protein was calculated using GraphPad Prism. (B) Blocking of murine HVEM-BTLA interaction: After the incubation with Anti-4CB1 mAb or isotype control (as described above, with 0.02-20 μg/mL), plate was incubated for two hours with PBS or 2.5 μg/mL of mouse BTLA-Fc tag protein followed by incubation for one hour with mouse BTLA/CD272 Biotin Antibody and incubation for 30 minutes with peroxidase-streptavidin and addition of TMB and stop solution. Absorbance was read at 450nM and 570nM.

| **Patient Number** | **Indication** |
| --- | --- |
| 1 | Colon |
| 2 | Hepatocellular |
| 3 | Colon |
| 4 | Ovarian |
| 5 | Endometrial |
| 6 | Endometrial |
| 7 | Colon |
| 8 | Ovarian |
| 9 | Melanoma |
| 10 | Renal |
| 11 | Bladder |
| 12 | Ovarian |
| 13 | Ovarian |
| 14 | Endometrial |
| 15 | Pancreatic |
| 16 | Renal |
| 17 | Renal |
| 18 | Pancreatic |
| 19 | Renal |
| 20 | Gastric |
| 21 | Endometrial |
| 22 | Lung |
| 23 | Colon |
| 24 | Lung |
| 25 | Renal |
| 26 | Bladder |
| 27 | Renal |
| 28 | Endometrial |
| 29 | Colon |
| 30 | Renal |
| 31 | Lung |
| 32 | Hepatocellular |
| 33 | Hepatocellular |
| 34 | Renal |
| 35 | Colon |
| 36 | Colon |
| 37 | Renal |
| 38 | Ovarian |
| 39 | Melanoma |
| 40 | Renal |
| 41 | Hepatocellular |
| 42 | Renal |
| 43 | Colon |
| 44 | Melanoma |
| 45 | Melanoma |
| 46 | Hepatocellular |
| 47 | Colon |
| 48 | Endometrial |
| 49 | Endometrial |

**Table S1. *Ex-vivo* cancer samples obtained from fresh human tumor tissues.**

| **Indication** | **Anti-4CB1 response rate** | | **Anti-PD1 response rate** | | **Overlap of responding samples** |
| --- | --- | --- | --- | --- | --- |
| Bladder | 0% | (0/2) | 0% | (0/2) | 0 |
| Colon | 44% | (4/9) | 11% | (1/9) | 1 |
| Endometrial | 29% | (2/7) | 29% | (2/7) | 1 |
| Gastric | 0% | (0/1) | 0% | (0/1) | 0 |
| Hepatocellular | 40% | (2/5) | 40% | (2/5) | 1 |
| Lung | 0% | (0/3) | 0% | (0/3) | 0 |
| Melanoma | 75% | (3/4) | 25% | (1/4) | 1 |
| Ovarian | 20% | (1/5) | 0% | (0/5) | 0 |
| Pancreatic | 0% | (0/2) | 0% | (0/2) | 0 |
| Renal | 18% | (2/11) | 9% | (1/11) | 1 |

**Table S2. *Ex-vivo* cytotoxicity experiments response rate per indication**. Results are shown as percent and ratio in parentheses of samples with increased cytotoxicity out of total number of samples per indication. Overlap of responding samples represent the number of samples showing increased cytotoxicity with both Anti-4CB1 and Anti-PD1.

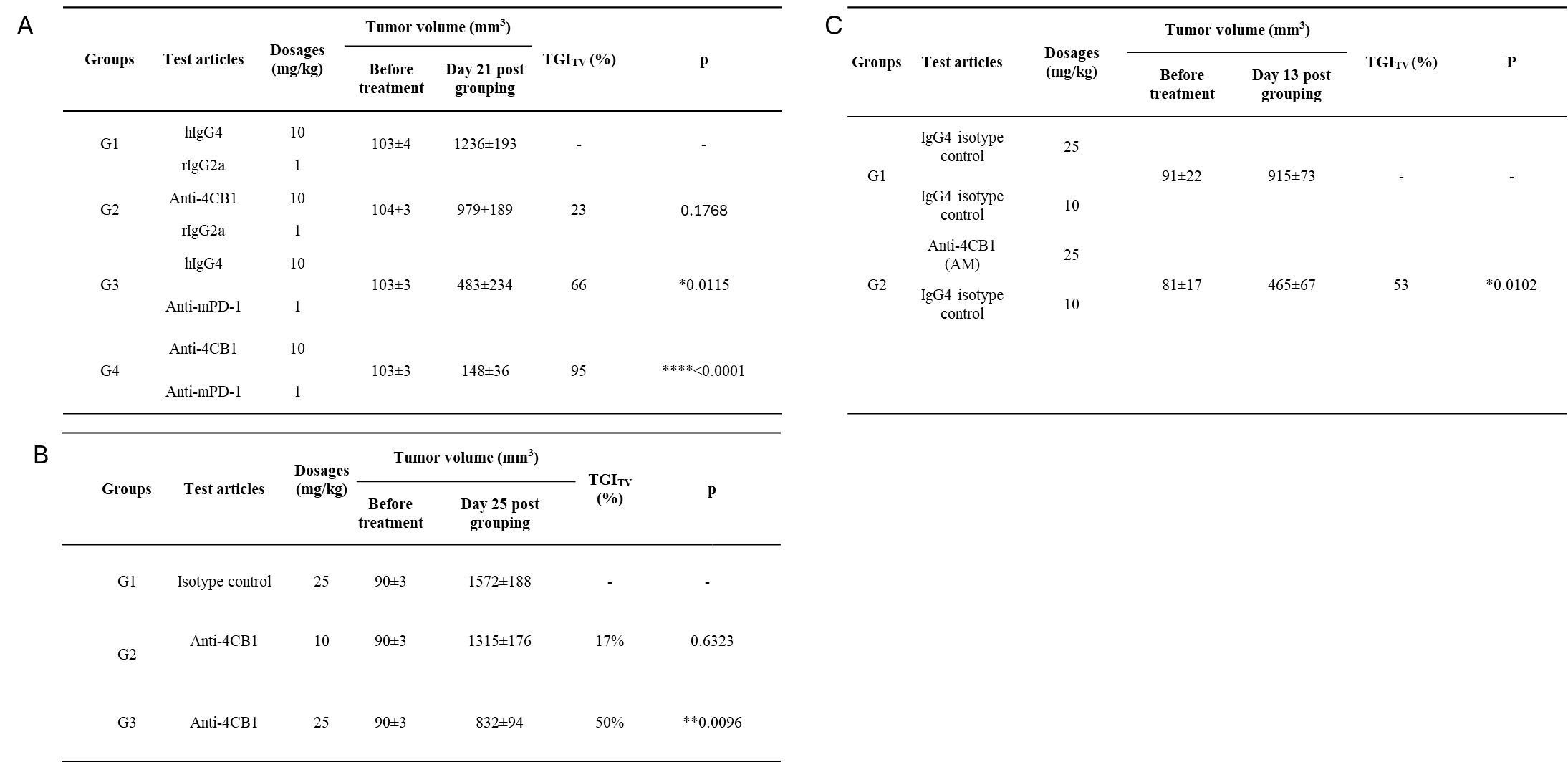

**Table S3. Tumor volume upon antibody treatment in *in-vivo* models**

(A) Tumor volume upon antibody treatment in B-hBTLA mice inoculated with B-hHVEM MC38 cells. Measures of tumor growth showed as ratio of size of the tumor before treatment and on day 21 post first dosing. (B) Tumor volume upon antibody treatment in B-hBTLA/hHVEM mice inoculated with B-hHVEM MC38 cells. Measures of tumor growth showed as ratio of size of the tumor before treatment and on day 25 post first dosing. (C) Tumor volume upon treatment in myeloid-boosted humanized NCG mice inoculated with CTG-1769 melanoma model. Measures of tumor growth showed as ratio of size of the tumor before treatment and on day 13 post first dosing.
